# Su*p4h5*-L19 activation tagging line partially restores root hair growth in *p4h5* mutant by introducing small transcriptomic changes in *Arabidopsis thaliana*

**DOI:** 10.1101/2024.06.14.599040

**Authors:** Tomás Urzúa Lehuedé, Gerardo Núñez-Lillo, Juan Salgado Salter, Romina Acha, Miguel Angel Ibeas, Claudio Meneses, José M. Estevez

## Abstract

An specific group of 2-oxoglutarate (2OG) dioxygenases named as Prolyl 4-Hydroxylases (P4H) produce trans-4-hydroxyproline (Hyp/O) from peptidyl-proline, catalyzing proline hydroxylation of cell wall glycoproteins EXT, AGPs, and HRGPs in plant cells, a crucial modification for *O*-glycosylation. Out of the Arabidopsis thaliana 13 P4Hs, P4H5 regulate root hair cell elongation and T-DNA insertional *p4h5* mutant has arrested cell elongation and shortened root hairs. P4H5 selectively hydroxylates EXT proline units indicating that EXT proline hydroxylation as an essential modification for root hair growth. In this work, we isolate an activation-tagging line called Su*p4h5*-L19/*p4h5* (*p4h5*-L19) that partially suppressed root hair phenotype in the *p4h5* mutant background. The T-DNA insertion site was mapped by Thermal Asymmetric Interlaced PCR (TAIL-PCR) followed by PCR product sequencing and the T-DNA is inserted at the beginning of the sixth exon of the AT3G17750 gene, an uncharacterized cytosolic kinase. By analyzing expression changes and mutants analysis in this loci, no clear direct effect was detected. By RNA-seq analysis, it become clear that *p4h5*-L19 may largely reverse the genetic alterations caused by the *p4h5* mutant in Wt Col-0, particularly at 10°C where there is an increase in root hair growth, with a total of 14 genes that have been activated and 83 genes that have been suppressed due to the enhancer of the activation tagging L19 in *p4h5* L19 compared to *p4h5*. Among these genes, 3 of them, Tonoplast Intrinsic Proteins (TIPs), were identified to be root hair specific (TIP1;1, TIP2;2, and TIP2;3) and the corresponding mutants for two of them (TIP1;1 and TIP2;3) showed reduced root hair growth response at low temperature. This study unmasked new components of the root hair growth response at low temperature that works independently of the *O*-glycosyated EXTs in the cell walls.

## 1. Introduction

P4Hs are 2-oxoglutarate (2OG) dioxygenases (EC 1.14.11.2) that generates trans-4-hydroxyproline (Hyp/O) from peptidyl-proline with Fe^2+^, O_2_, and ascorbate cofactors. In plant cells, P4Hs only have the catalytic α-subunit (Tiainen et al. 2005; Koski et al., 2007; Koski et al., 2009) and catalyzes the proline hydroxylation of EXT, AGPs and related Hydroxyproline Rich-GlycoProteins (HRGPs) (Borassi et al. 2016; Marzol et al. 2018), key modification for the subsequent *O*-glycosylation. The growth of root hairs was hindered when treated with P4H inhibitors DP (a,a-dipyridyl) and EDHB (ethyl-3,4-dihydroxybenzoate), as they blocked the hydroxylation of peptidyl-proline HRGP and significantly inhibited cell elongation. This indicates a direct connection between proline hydroxylation and the growth of root hairs (Velasquez et al., 2011, 2015a,b). *Arabidopsis thaliana encodes* 13 P4Hs (Hieta and Myllyharju, 2002; Tiainen et al., 2005; Velasquez et al., 2011; Velasquez, 2015a,b) and P4H2, P4H5, and P4H13 were shown to be highly expressed in root epidermal trichoblast cells and affect root hair growth (Velasquez et al., 2015a,b). In agreement, T-DNA insertional mutants for P4H2, P4H5, and P4H13 displayed cell elongation arrest and shortened root hair morphologies (Velasquez et al. 2011). The *p4h5* mutant has the most abnormal cell wall structure, reflecting a similar phenotype than the triple mutant *p4h2 p4h5 p4h13* (Velasquez et al., 2015a,b). In contrast, P4H5 overexpression causes over-elongated root hair (Velasquez et al. 2011; 2015a,b). P4H5 selectively hydroxylates three of EXT’s first four proline units (SOOOP) in a particular order (Velasquez et al., 2015b) and together this suggests that in root hair cells, P4H5 initiates and maintains proline hydroxylation of EXTs and related glycoproteins as essential modification for *O*-glycosylation linked to root hair cell growth. The expression of P4H::GFP fusions, controlled by their endogenous promoters, showed that P4H2, P4H5, and P4H13 are mostly expressed in root epidermal trichoblast cells and developing root hairs. Additionally, these proteins are found in the endoplasmic reticulum (ER) and Golgi compartments (Velasquez et al., 2011, 2015b). These findings indicate that the process of proline hydroxylation of HRGPs may start in the endoplasmic reticulum (ER) and be completed in the Golgi apparatus. Once secreted, monomeric extracellular EXTs have a rod-like shape with a polyproline-II helical conformation and these structures are made more stable by the presence of Hyp-O-glycans (Stafstrom and Staehelin in 1986; Owens et al. in 2010; Velasquez et al. in 2011; Velasquez et al. 2015b). Furthermore, several EXTs undergo crosslinking and insolubilization in the plant cell wall via Tyr-based motifs, in addition to EXT *O*-glycosylation (Lamport et al., 2011). Secreted type-III PERs are believed to aid in the formation of both intramolecular and intermolecular covalent Tyr-Tyr crosslinks by producing *iso-*dityrosine units and pulcherosine or di-*iso-*dityrosine, respectively (Brady et al., 1996, 1998). However, the exact molecular mechanisms responsible for this process have not yet been fully understood.

In several cases, examination of loss-of-function mutations in some potential genes does not provide particular gene targets for functional characterization in a particular biological process. On the contrary, we conducted a gain-of-function, forward genetic screen utilizing an activation tagged population of Arabidopsis. One of our goals for doing activation-tagging screens is to find genes that have overlapping functions and that are not readily recognized by mutations that cause loss of function. Activation-tagged mutants have successfully been used to identify new genes in specific signaling pathways (Grant et al. 2003; Aboul-Soud et al. 2009; Xia et al. 2004; Yadeta et al. 2011; Xiao and Anderson, 2015). Activation tagging is a process where regulatory sequences are randomly inserted into a plant genome using T-DNA or transposons (Weigel et al., 2000). Gain-of-function mutants arise from the activation of genes near the integrations, leading to increased transcriptional activity. In rare cases, the activation tagging may also induce transcriptional changes in a longer range of 10 or more kb (Lewin 2008). The purpose of this screen was to directly discover mutants that are able to rescue the strong root hair phenotype of the *p4h5* mutant, with defects in the cell wall structure. A single line named Sup4h5-L19 with a partially rescue phenotype was identified. Surprisingly, this characteristic was not attributed to the activation of any of the genes close to the insertion sites nor due to the interruption of the gene where the T-DNA was located.

Instead, we found small global transcriptional changes of hundreds of genes were restored in the Su*p4h5* L19 in comparison to *p4h5* mutant and much more similar to Wt Col-0.

## 2. Results

To identify new genes involved in the regulation of root hair growth linked to hydroxylation and concomitant *O*-glycosylation pathway of EXTs, an activation-tagging screening was performed on the background of *p4h5* mutant which has much shorter root hairs than Wt Col-0. Plants of *p4h5* were transformed with *Agrobacterium tumefaciens* carrying the activation tagging vector pSKI015, which contains four copies of the CaMV virus enhancer 35S (Weigel et al. 2000). Transformed *p4h5* plants were selected using the Basta selection marker and the population generated of 1,000 plants were analyzed looking for a line in which root hair growth had been restored similar to Wt Col-0 levels. The best 50 candidates were analyzed and a line called Su*p4h5*-L19/*p4h5* (*p4h5*-L19) for the suppressor of *p4h5* was obtained that partially rescued the root hair phenotype of *p4h5* by 50%. The pSKI105 T-DNA insertion site was mapped by Thermal Asymmetric Interlaced PCR (TAIL-PCR) followed by PCR product sequencing and the T-DNA is inserted at the beginning of the sixth exon of the AT3G17750 gene, an uncharacterized cytosolic kinase, affecting its reading frame (**Figure 1A**). A SALK mutant line of AT3G17750 (Salk_064507) was obtained and crossed with *p4h5*, leading both mutations to an homozygous state and named as *p4h5* SALK-L19 (**Figure 1B**). The root hair phenotype was indistinguishable from that of *p4h5*, which suggests that the effect observed in *p4h5*-L19 is not due to the disruption of AT3G17750 gene expression but to changes in the expression of neighboring genes by the activation tagging construct. In agreement, homozygous SALK_064507 in Wt Col-0 background (WT SALK-L19) did not show any distinguishable root hair phenotype from Wt Col-0 (**Figure 1C**). In addition, L19 was crossed to Wt Col-0 (WT L19) and the effect of disruption on AT3G17750 on Wt Col-0 background did not show any abnormal root hair phenotype when compared to Wt Col-0 (**Figure 1C**). Based on these results, it is clear that the rescue of the root hair phenotype in *p4h5* L19 is not due to the T-DNA insertion on AT3G17750.

**Figure 1.**
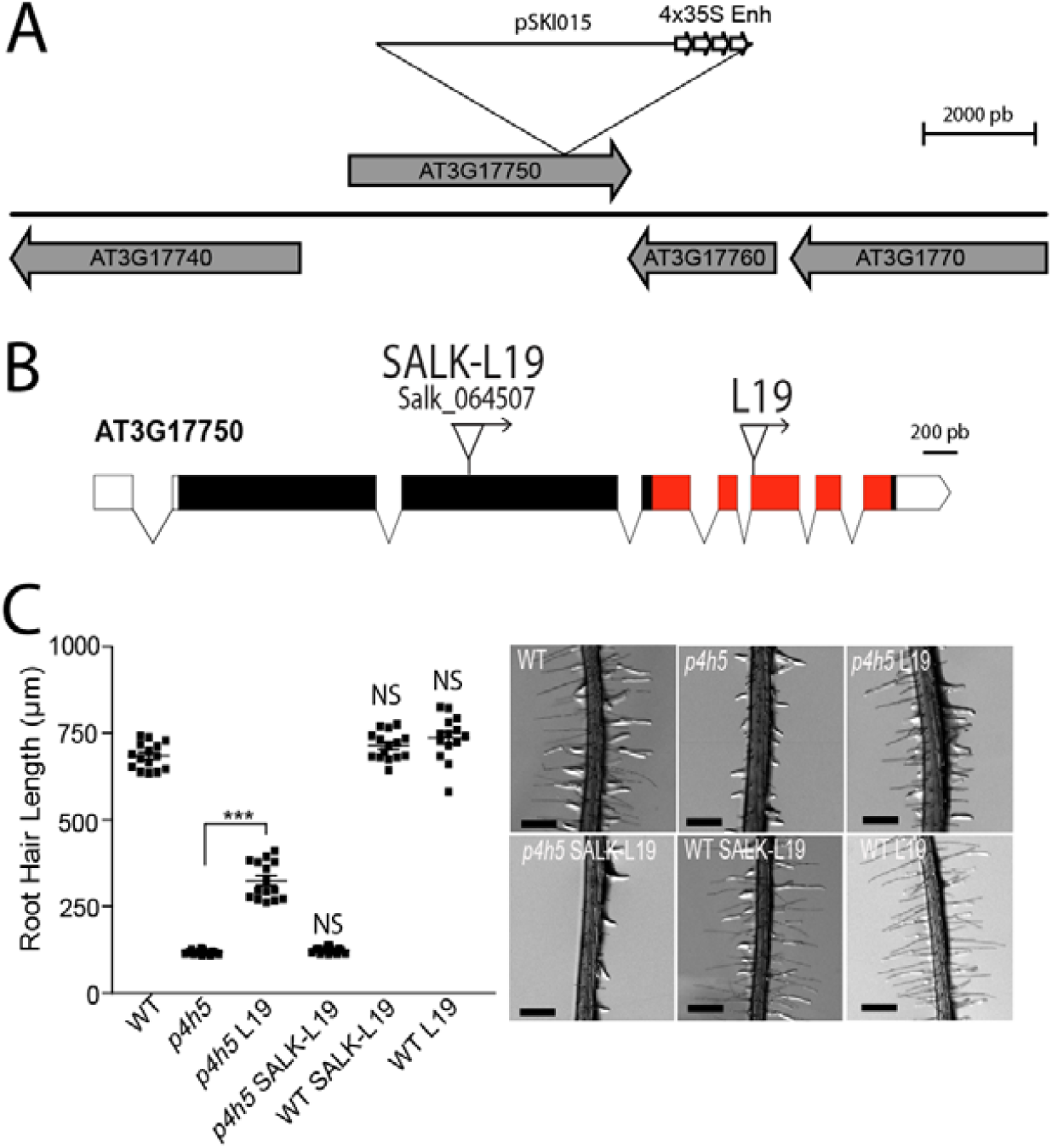
Partial rescue of *p4h5* root hair phenotype by activation tagging SuP4H5-L19 (L19). **(A)** Place of insertion of T-DNA corresponding to Activation Tagging pSKI015 plasma in the suppressor line *p4h5*-L19 (L19). It contains 4 copies of the enhancer 35S (4×35S Enh). **(B)** Detail of the structure of the gene AT3G17750 indicating the place of introduction of the T-DNA of *p4h5*-L19 and the *p4h5* SALK-L19 (Salk_064507). The boxes represent the exons and the connector lines, the introns. The coding sequence is shown in black and red and the untranslated regions 5’ and 3’ are shown in white. The region in red color indicates the kinase domain. (**C**) Quantification of root hair length in roots grown at 22°C in *p4h5, p4h5* L19, *p4h5* SALK-L19, WT SALK-L19 and WT L19. The data were analyzed using ANOVA of one factor followed by Bonferroni post-hoc comparisons, (***) P=0.001. On the right, representative photos of the root hair phenotype. Scale Bar = 700 μm.

To test if local or more distant changes in gene expression are triggered by the effect of *p4h5*-L19, we performed an RNA-seq analysis of 24hs treated root at 10°C (**Figure 2A**) where root hair growth is triggered by two folds by the reduction in mobility and accessibility of specific nutrients in the media (Moison et al. 2021, Pacheco et al. 2021, Pacheco et al. 2023a,b). First, we tested the root hair phenotype *p4h5*-L19 at 10°C and a similar degree of partial rescue of *p4h5* is observed although root hair growth is enhanced proportionally in all lines (**Figure 2B**). Since activation tag enhancer elements can potentially enhance the expression of multiple genes in their vicinity (Weigel et al., 2000; Jeong et al. 2002), we test if *p4h5*-L19 might trigger expression changes in genes close to the AT3G17750 insertional site. To establish if the activation tagging of *p4h5*-L19 is affecting changes in the surrounding genetic environment close to AT3G17750, the expression levels of surround genes were assessed in Wt Col-0, *p4h5* and *p4h5*-L19 at 22°C and at 10°C (**Figure 2C**). The insertion L19 was clearly found to have an strong effect on the expression of AT3G17750 (**Figure 2C**) but no gene expression changes were detected in AT3G17740 and AT3G17770 around L19 (**Figure 2C**). Based on these results, *p4h5*-L19 is able to partially rescue *p4h5* abnormal short root hair phenotype but this is not mediated by the insertion in AT3G17750 or in any detectable changes in the expression of the nearby genes in this genomic region of 10-50 kb.

**Figure 2.**
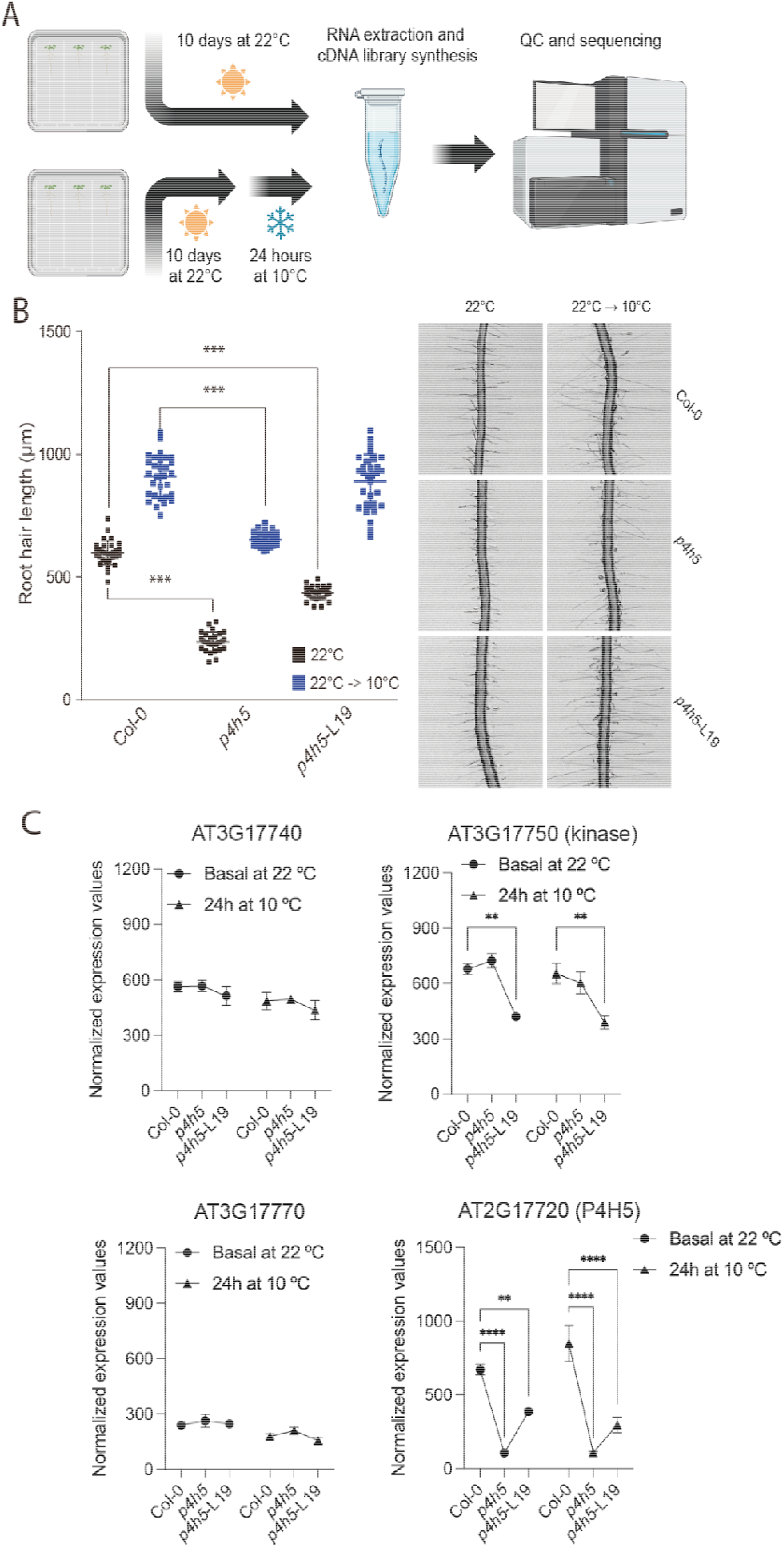
The activation tagging Sup4h5 L19 does not change expression around the AT3G177750 locus. **(A)** RNA seq approach to study changes in gene expression in Wt Col-0, *p4h5* and *p4h5* L19 at 22°C and at 10°C for 24hs. (**B**) Quantification of root hair length in *p4h5, p4h5-L19* and Wt Col-0 at 22°C and at 10°C for 24hs. The data were analyzed using ANOVA of one factor followed by Bonferroni post-hoc comparisons, (***) P=0.001. On the right, representative pictures of the root hair phenotypes. Scale Bar = 700 μm. (**C**) Gene expression levels assessed by RNA-seq reads of the genes (AT3G17740 and AT3G17770) close to the insertion site of the activation tagging suppressor line *p4h5*-L19 (AT3G17750) including P4H5.

The activation tagging lines might have a long range effect on the expression of neighboring genes and it can influence the expression of genes up to several kilobases away, occasionally more than 10 kb from the enhancer insertion (Lewin 2008). Based on the global expression analysis, 209 differentially expressed genes (DEGs) were found in the comparisons Col-0, *p4h5* and *p4h5*-L19 grown at 22°C for 10 days and those growth at 10 days at 22°C and 24hs at 10°C (**Figure 3A**). 209 DEGs identified are shown in the Heatmap, where 9 clusters are defined from cluster 1 located at the top and cluster 9 is the one at the bottom (**Figure 3A**). Partial Least Squares regression-Discriminant Analysis (PLS-DA) of component 1 and 2 first, where we observe the separation between basal treatment and 24hs of 10°C (component 1; 40%) and separation of Col-0 with the mutant samples (Component 2; 6%), we also observe there the segregation between *p4h5* and *p4h5*-L19 but only at 24h of low temperature treatment (**Figure 3B**). In the second PLS-DA, it is shown the component 5 (4%) that allows us to observe the separation between *p4h5* and L19 in the 22°C treatment. Interestingly, in Cluster 9 we can identify AT2G17720 gene (P4H5) and the AT3G17050 gene (kinase where L19 T-DNA insertion is located). GO analysis highlighted terms related to oxygen, hypoxia, nutrients, and energy processes (**Figure 3C**). It is clear from these global analyses that *p4h5*-L19 is able to restore to Wt Col-0 most of the genetic changes introduced by the *p4h5* mutant, specially at 10°C where root hair growth is enhanced. Finally, we identified 14 genes that are upregulated and 83 downregulated genes triggered by the enhancer of the activation tagging L19 in *p4h5* L19 versus *p4h5* (**Supplementary Table S1**). Then, we filtered those that are specifically and/or highly expressed in root hair cells previously identified by single cell RNA-seq studies (Brady et al., 2007; Denyer et al., 2019; Jean-Baptiste et al., 2019; Ryu et al., 2019; Shulse et al., 2019; Zhang et al., 2019). There are 9 dysregulated root hair genes (RHG) in *p4h5-*L19 and 3 are Tonoplast Intrinsic Proteins (TIP1;1, TIP2;2 and TIP2;3). TIP1;1 and TIP2;3 are highly expressed in root hair cells according to the single root cell RNA-seq database https://rootcellatlas.org/ (**Figure 4A**). T-DNA mutants for TIP1;1 and TIP2:3 were isolated (**Figure 4B**) and most of them showed a reduced response to low temperature growth, especially *tip1;1* (**Figure 4C**). This indicates that these three root hair specific TIP are important players in root hair growth under low temperature and their expression may be controlled, most likely indirectly, by the activation tagging L19.

**Figure 3.**
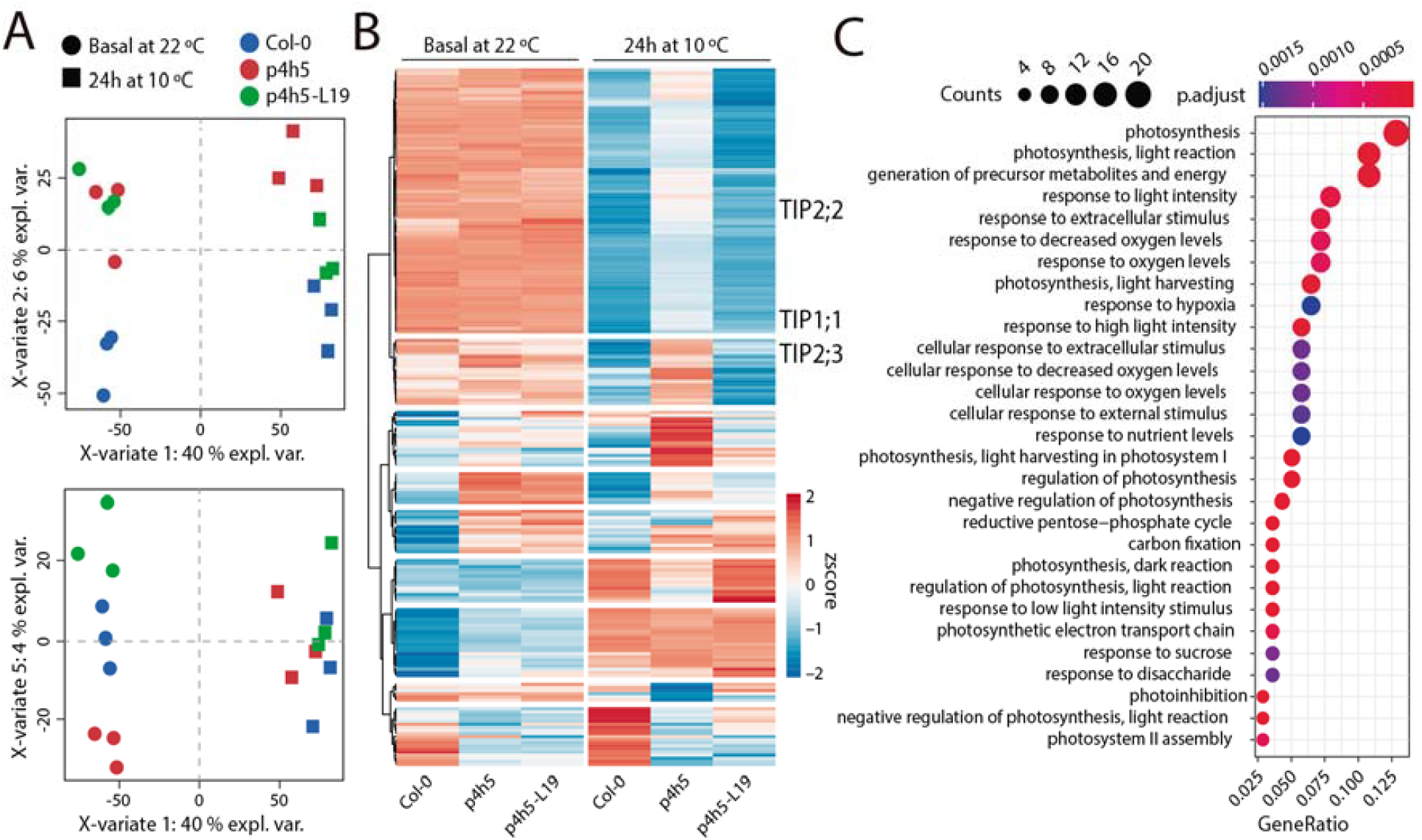
Global transcriptional changes of 209 DEG in *p4h5*-L19 triggered the partial rescue of *p4h5* to Wt Col-0 root hair phenotype. **(A)** Partial Least Squares Regression-Discriminant Analysis (PLS-DA) on component 1 and 2. In component 1 (40%), the separation between the basal treatment and 24 hours of 10°C. In component 2 (6%), the separation between the Col-0 samples and the mutant *p4h5* and *p4h5* L19. Additionally, the segregation between *p4h5* and *p4h5*-L19, but only at 24 hours of low temperature treatment is shown. The second PLS-DA analysis reveals that component 5 (4%) effectively demonstrates the distinction between p4h5 and L19 in the 22°C treatment. (**B**) Heat-map showing the hierarchical gene clustering for 209 differentially expressed genes (DEG) between room temperature growth (22°C) and low temperature (10°C) growth in wild type Col-0, *p4h5* mutant and *p4h5*-L19 roots. On the right, tree TIPs are indicated in Cluster 1 and Cluster 2. **(C)** Gene Ontology analysis results depicting the top most significantly 30 enriched GO terms are shown as bubble plots on the right. The size of the points reflects the amount of gene numbers enriched in the GO term. The color of the points means the *p*-value.

**Figure 4.**
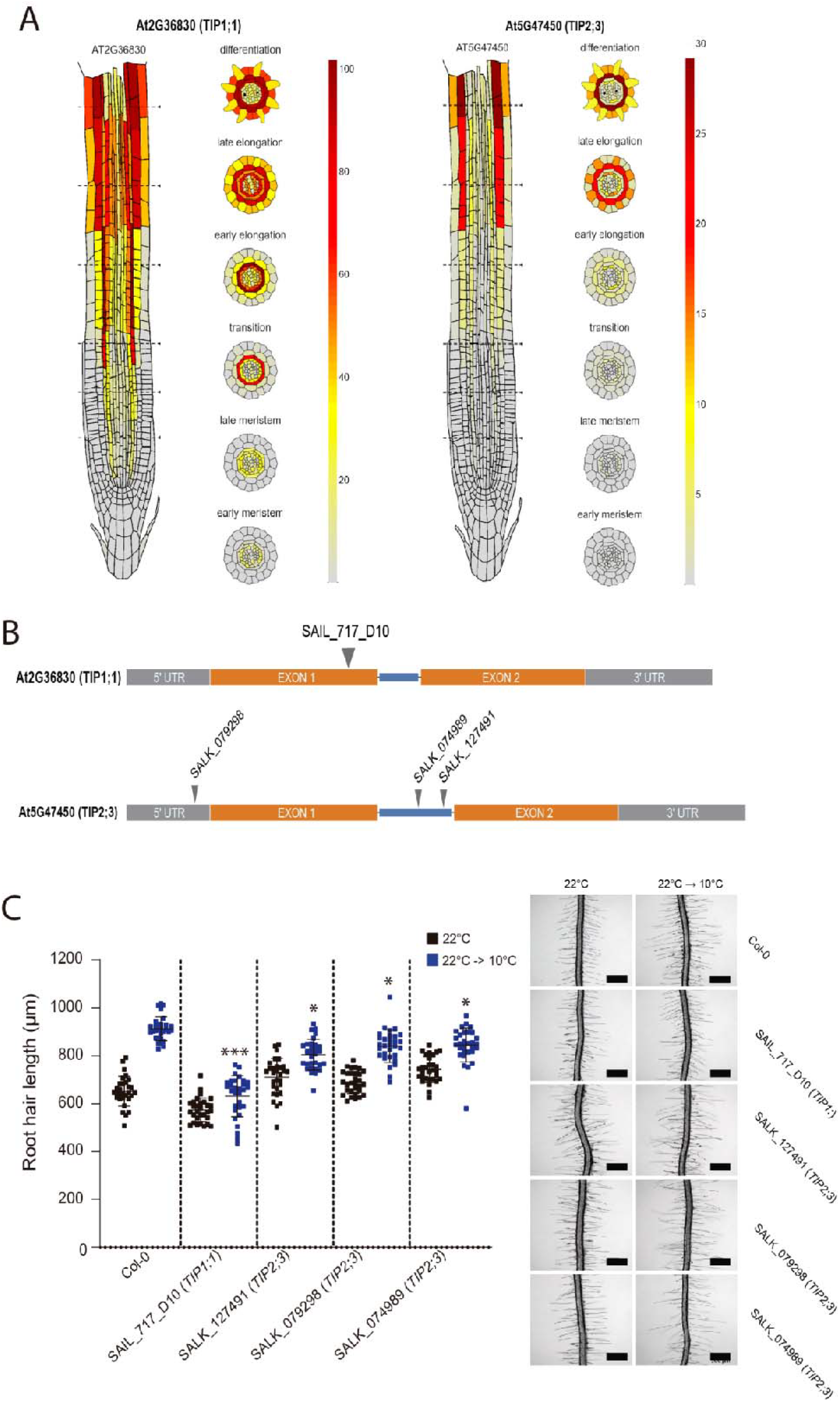
TIP1;1 and TIP2:3 are involved in the root hair growth response to low temperature. **(A)** Expression profile of TIP1;1 and TIP2;3 based on the single root cell RNA-seq dataset https://rootcellatlas.org/. (**B**) TIP1;1 and TIP2;3 gene structure and place of insertion of T-DNA for the mutants isolated. The boxes represent the exons and the blue connector lines, the introns. The coding sequence is shown in orange and the untranslated regions 5’ and 3’ are shown in grey. (**C**) Quantification of root hair length in Wt Col-0 and TIP mutants at 22°C and at 10°C for 3 days. The data were analyzed using ANOVA of one factor followed by Bonferroni post-hoc comparisons, (***) P=0.001 (*) P=0.01. On the right, representative pictures of the root hair phenotypes. Scale Bar = 500 μm.

## 3. Discussion

By using the activation tagging approach, a strong promoter sequence is randomly inserted into the plant genome and this promoter sequence drives the expression of nearby genes, often resulting in overexpression of those genes. By activating the expression of nearby genes, activation tagging can lead to the identification of genes involved in specific biological processes or traits of interest. In our case, we have identified the line *p4h5*-L19 that is able to partially suppress the abnormal *p4h5* root hair phenotype by introducing gene expression changes in a small group of 97 DEG, most of which reverted to wild Col-0 pattern. When filtering which are specifically expressed in root hair cells, we identified three TIPs (TIP1;1, TIP2;2 and TIP2;3). Root hair cells necessitate the permanent increment of the surface area of the cell wall and it must produce internal turgor pressure by absorbing water and several other molecules (Cosgrove 2023). Aquaporins, which are water channels, facilitate the passage of water and several other molecules such as urea, CO_2_, NH_3_, H_2_O_2_, and metalloids (Liu et al., 2003; Uehlein et al., 2003; Jahn et al., 2004; Bienert et al., 2007; Bienert et al., 2008; Bienert et al., 2011; Bienert et al., 2014) across cell membranes to maintain turgor during cell wall expansion. Our analysis revealed three TIPs aquaporin genes were dysregulated in *p4h5 L19* (**Supplementary Table S1**). TIP2;2 regulates water and solutes in tonoplast (Uenishi et al 2014) and it is involved in response to abiotic stress through the modulation of physiological components (Feng et al. 2018) and TIP2;3 facilitates NH_3_ into the vacuole (Loque et al. 2005). This suggests that these proteins could have a role in maintaining a greater turgor pressure, which in turn facilitates proper cell elongation. In agreement, it was shown that other three Arabidopsis TIP isoforms (TIP1;1, TIP1;2, and TIP2;1) play a crucial role in regulating cellular water transport throughout the process of lateral root primordia morphogenesis and emergence also related to cell elongation (Reinhardt et al. 2016). These 3 TIPs identified here represent an independent pathway activated in *p4h5-*L19 that rescues the severe defect on hydroxylation and *O*-glycosylation of EXTs and related HRGPs in the *p4h5* mutant. This highlights the possibility that the plant cell, in this case root hairs, fine-tunes the turgor pressure at the vacuole level to overcome structural changes at the cell walls. This molecular connection was previously shown for the *Catharanthus roseus* receptor-like kinase-1-like kinase (CrRLK1L) FERONIA (FER) and its interactors, the *O*-glycosylated cell wall components of the leucine-rich repeat extensins (LRXs) type and the oscillating pectin methyl esterified linked to the vacuole expansion in root epidermal cells (Feng et al. 2018; Dunser et al. 2019; Herger et al. 2019; Rößling et al. 2024).

## 4. Conclusions

Overall, we have found an activation tagging line *p4h5*-L19 that is able to revert small global transcriptomic changes in the mutant *p4h5* and partially rescue the abnormal root hair phenotype. The three discovered TIPs described here correspond to an autonomous route that is activated in *p4h5-*L19, which effectively corrects the significant impairment in hydroxylation and *O*-glycosylation of EXTs and associated HRGPs in *p4h5*. This emphasizes the potential for the plant cells to precisely adjust the turgor pressure at the vacuole level in order to counteract changes in the structure of the cell wall. A putative new root hair growth pathway composed by three TIPs that counteracts defective cell wall HRGPs *O*-glycosylation in *p4h5* deserves to be explored in the future.

## 5. Material and methods

### 5.1. Plant Material and Growth Conditions

All the *Arabidopsis thaliana* lines used were in the Columbia-0 (Col-0) background. Seedlings were surface sterilized and stratified in darkness at 4°C for 3 days before being germinated on ½ strength 0,5X MS-MES (Duchefa, Netherlands) on 0.8% Plant Agar™ (Duchefa, Netherlands) on 120 x 120 mm square petri dishes (Deltalab, Spain) in a plant growth chamber in continuous light (120 μmol s^−1^ m^−2^). Root hair phenotype characterization. seeds were surface sterilized and stratified in darkness for 3 days at 4 °C. Then grown on ½ strength MS agar plates, in a plant growth chamber at 22 °C in continuous light (120 μmol s^−1^ m^−2^) for 7 days at 22°C as a pretreatment and then at 10°C (low temperature treatment) for 3 days. Measurements were made in 10 days. For quantitative analysis of RH phenotype, 10 fully elongated RHs were measured (using the ImageJ software) from each root (n=20) grown on vertical agar plates. After treatment only new RH was measured. Images were captured with a Leica EZ4 HDStereo microscope (Leica, Germany) equipped with the LAZ ez software. Results were expressed as the mean ± SD using the GraphPad Prism 8.0.1 (USA) statistical analysis software.

### 5.2. Plant genotyping

For the identification of T-DNA knockout lines, genomic DNA was extracted from rosette leaves. Confirmation by PCR of a single and multiple T-DNA insertions in the genes were performed using an insertion-specific LBb1.3 (for SALK lines) primer in addition to one gene-specific primer. In this way, we isolated homozygous for all the genes. Arabidopsis T-DNA insertions lines (*tip1:1* SAIL_717_D10, *tip2;3* SALK_079298, *tip2;3* SAIL_074989, *tip2;3* SALK_127491) were obtained from Arabidopsis Biological Resource Center (ABRC, https://abrc.osu.edu/). Using standard procedures homozygous mutant plants were identified by PCR genotyping with the gene specific primers listed in *Supplementary Table S2*. T-DNA insertion sites were confirmed by sequencing using the same primers.

### 5.3. Plant transformation and transgenic plant selection

1,000 *p4h5* mutant plants that were about 5 weeks old were transformed with the activation tagging vector pSKI015 (Weigel et al., 2000) construct in *Agrobacterium tumefaciens* by the floral dip method (Clough and Bent, 1998). T1 seeds were selected on soil containing Basta and seeds of each resistant plant line were collected. Root hair of the transgenic plants T2 were assessed as described above, and confirmed in subsequent generations. The best 50 transgenic lines isolated as revertant of *p4h5* root hair phenotype, only *p4h5-*L19 was followed to further characterization.

### 5.4. Thermal Asymmetric Interlaced PCR (TAIL-PCR) of L19

The *p4h5* mutant plants were transformed with the activation tagging vector pSKI015 (Weigel et al., 2000). The *p4h5* L19 dominant mutant was isolated from approximately 10,000 mutant plants. Genomic DNA was isolated from 10-day-old L19 mutant seedlings by using a DNeasy Plant Mini Kit (Qiagen, www.qiagen.com) according to the manufacturer’s instructions. TAIL-PCR was performed as described by Liu et al. (1995). The left border of the T-DNA border-specific primer used in the first and second round TAIL-PCR cycling is LB3 (TTGACCATCATACTCATTGCTG). The degenerate primer pools AD1 (WGCNAGTNAGWANAAG) and AD2 (AWGCANGNCWGANATA) (Liu et al., 1995) were used per round of TAIL-PCR cycling. The PCR product was then purified by QIAquick PCR Purification Kit (QUIAGEN) and it was sequenced using LB3 primer. The product was then cloned in pGEM-T and sequenced.

### 5.5. RNA-seq analysis

For the RNA-seq analysis, seedlings were grown on ½ strength MS agar plates, in a plant growth chamber at 22 °C in continuous light (120 μmol s^−1^ m^−2^) for 10 days at 22°C as a pretreatment and then at 10°C (moderate-low temperature treatment) for 24hs. We analyzed a dataset with 6 factor groups (two time points and three genotypes: Col-0 0hs 22°C, Col-0 24hs 10°C, *p4h5* 0hs 22°C, *p4h5* 24hs 10°C, SuP4H5-L19 0hs 22°C, SuP4H5-L19 24hs 10°C) each with three biological replicates giving 18 samples in total. Total RNA was extracted from 30 mg of frozen root tissue. Frozen root samples were ground in liquid nitrogen and total RNAs were extracted using E.Z.N.A Total RNA Kit I (Omega Bio-tek, Georgia, USA). RNA quantity and purity were evaluated with a Qubit®2.0 fluorometer (InvitrogenTM, Carlsbad, CA, USA) using a QubitTM RNA BR assay kit. RNA integrity and concentration were assessed by capillary electrophoresis using an automated CE Fragment AnalyzerTM system (Agilent Technologies, Santa Clara, CA, USA) with the RNA kit DNF-471-0500 (15nt). Total RNA-seq libraries were prepared according to the TruSeq Stranded Total RNA Kit (Illumina, San Diego, CA, USA) following the manufacturer’s instructions. Finally, the constructed libraries were sequenced using Macrogen sequencing services (Seoul, Korea) in paired end mode on a HiSeq4000 sequencer. For total RNA differential expression analysis, a quality check was performed with FASTQC software (Andrews, 2010). Then, the adapter sequences were removed, reads with a quality score less than 30 and length less than 60 nucleotides were eliminated using Flexbar (Dodt et al., 2012). Resulting filtered reads were aligned against Arabidopsis thaliana Araport 11 genome with the STAR aligner software. A total of 18 RNA libraries were sequenced, obtaining an average of 71,013,704 reads for each one, with a minimum and maximum value of 53,419,520 and 84,351,800 reads, respectively.

After filtering them by quality and removing adapters, an average of 97.7% of the reads remained and after aligning them against the *Arabidopsis thaliana* reference genome, between 96.0% and 98.8% of total reads were correctly aligned. For each library, the feature Counts software from the Rsubread package (Liao et al., 2019) was applied to assign expression values to each uniquely aligned fragment. Differential gene expression analysis was performed using the Bioconductor R edgeR package (Robinson et al., 2010). Differentially expressed genes (DEG) were selected with an FDR < 0.05 and a FC > |0.5|. To search for genetic functions and pathways overrepresented in the DEG lists, genetic enrichment analysis was performed using the Genetic Ontology (GO) database with the R package ClusterProfiler v4.0.5 (Yu et al., 2012), using the compareCluster function. The parameters used for this analysis were: lists of differentially expressed genes for each comparison in ENTREZID, enrichGO sub-function, the universe from the total of differentially expressed genes that present annotation as genetic background, Benjamini-Hochberg statistical test and a filter of FDR less than 0.05. Subsequently, the semantics filter of GO terms was performed using the simplify function of the same package using a p-value and q-value cutoff less than 0.05.

## Acknowledgements

We thank ABRC (Ohio State University) for providing T-DNA seed lines. J.M.E. is investigators of the National Research Council (CONICET) from Argentina. M.I. is supported by ANID FONDECYT POSTDOCTORADO [grant 3220138]. This work was supported by grants from ANPCyT (PICT2019-0015 and PICT2021-0514), by ANID – Programa Iniciativa Científica Milenio ICN17_022, NCN2021_010 and Fondo Nacional de Desarrollo Científico y Tecnológico [1200010] to J.M.E.

## Authors contributions

T..U. designed research and performed experiments. G.N.L. Performed the bioinformatic analysis. J.S.S. performed the activation tagging mapping. R.A. performed the mutant isolation. M.A.I. analyzed data. C.M. analyzed the bioinformatic data. J.M.E.designed experiments, analyzed data and wrote the manuscript.

**Supplementary Table S1.**
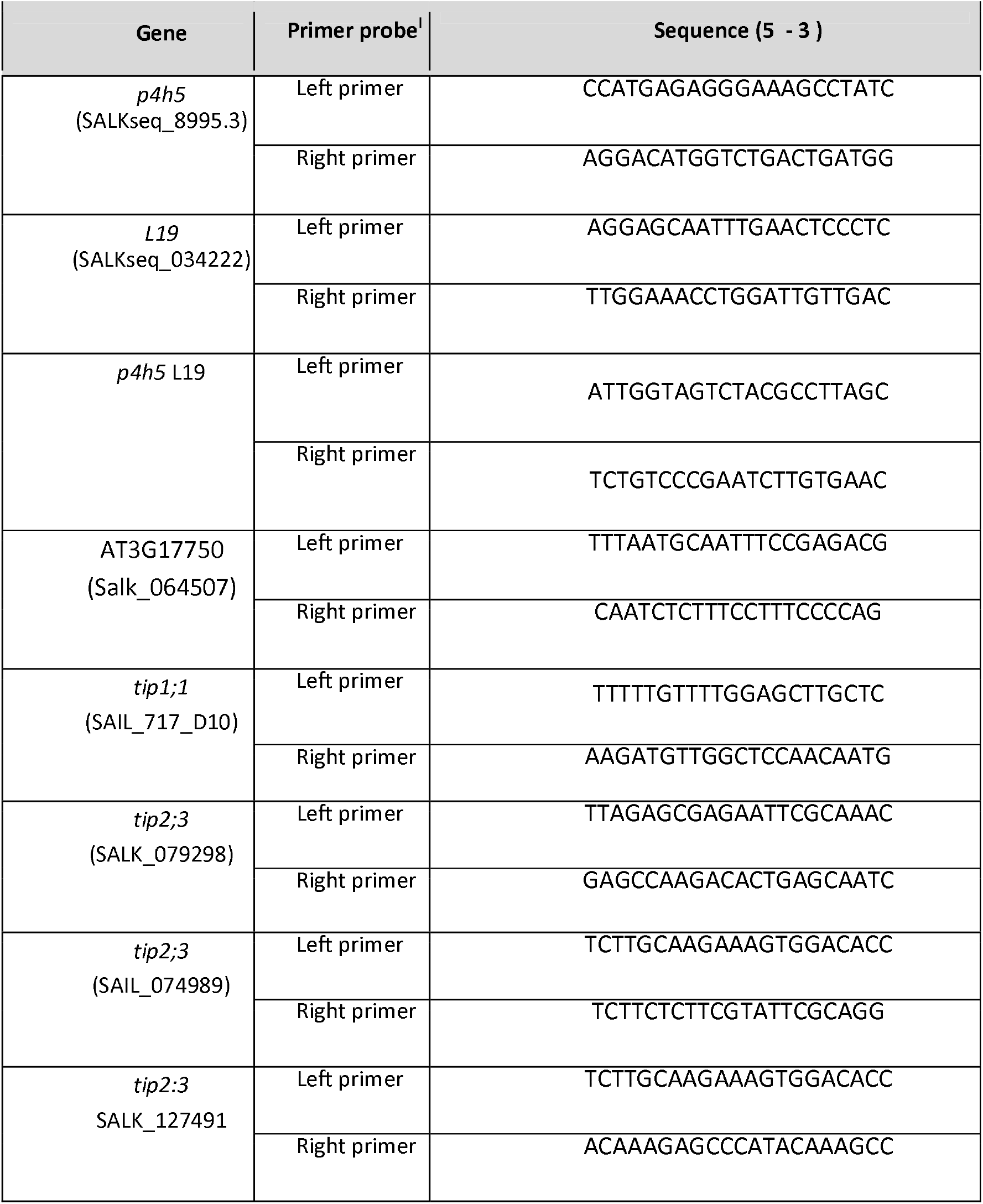
Oligonucleotide list. Primer sequences are shown next to their corresponding genes.

## Notes

### Competing Interest Statement

The authors have declared no competing interest.

